# *Wolbachia* manipulates host pre-imaginal learning in a parasitoid wasp

**DOI:** 10.1101/825455

**Authors:** Pouria Abroon, Ahmad Ashori, Anne Duplouy, Hossein Kishani Farahani

## Abstract

The Hopkin’s host-selection principle (HHSP) suggests that organisms at higher trophic levels demonstrate a preference for the host species on which they developed during larval stage. Although investigated in many herbivorous and predatory insects, the HHSP has, to our knowledge, never been tested in the context of insects hosting selfish endosymbiotic passengers such as the maternally inherited bacterium *Wolbachia pipientis*. Here, we investigate the effect of *Wolbachia* infection on host pre-imaginal learning in the parasitoid wasp *Trichogramma brassicae* (Hymenoptera: Trichogrammatidae). We compare host-choice in *Wolbachia*-infected and uninfected adult female parasitoids after rearing them on two different Lepidopteran hosts, namely the flour moth *Ephestia kuehniella* Zeller (Lepidoptera: Pyralidae) or the grain moth *Sitotroga cerealella* (Lep.: Gelechiidae). We show that in *T. brassicae, Wolbachia* affect the pre-imaginal learning ability of female wasps. *Wolbachia* infected wasps do not show any host preference and easily switch hosts in the laboratory, while uninfected wasps significantly prefer to lay eggs on the host species they developed on. We discuss how the facilitation of a generalist strategy by *Wolbachia* may allow *T. brassicae* to escape intraspecific competition with their uninfected counterparts, and may have important evolutionary consequences for the host and its symbionts.

## Introduction

Learning from experience is a property embedded into the survival strategies of most animals, including insects [1-3]. For example, learning from olfactory cues enables many insects to optimize foraging, mating, and the completion of other behaviors in their complex local environments [4-6]. Parasitoid wasps especially rely on such innate mechanisms to directly identify, detect and parasitize their hosts [7, 8], or indirectly detect the host plants or any other characteristics of the environment in which the parasitoid has previously encountered its host [9]. Consequently, olfactory learning in parasitoids may contribute to behavioral optimization through increasing the speed and the efficiency of detecting suitable hosts, which should also positively affect the fitness of the parasitoid [10-12]. In the Ichneumonoid parasitoid wasps, *Hyposoter horticola* and *Venturia canescens*, females use olfactory cues from conspecific deterrent markings, to avoid costly interference and competition for resources (hosts) between individuals [13, 14]. Similarly, it has been shown that *Trichogramma brassicae* wasps reared from larvae feeding on tomato, later parasitized significantly more hosts on tomato plants than on lettuce [15].

The learning process of olfactory cues in insects was previously suggested to either occur during adult emergence, or at the young adult stage [9, 16-19]. Learning at adult emergence relies on traces of chemical cues from hosts, inside or outside the host body, which influence adult behavior potentially during a ‘sensitive period’ associated with adult emergence [20-24] This phenomenon, learning at emergence, has been proposed to occur in two different ways. First, as a result of larval experiences, that implies that the larva can learn from its environment and that this memory can be transferred from pre-imaginal stages to the adult—the so-called Hopkins’ host selection principle [25]; second, at eclosion of the imago, in which the larval environment is carried over to the adult stages and olfactory learning occurs during the contact of the young wasp with olfactory cues at emergence—the so-called “chemical legacy” hypothesis [19, 26]. Two ectoparasitoid wasps, *Hyssopus pallidus* and *Aphidius ervi*, of the codling moth, *Cydia pomonella*, rely on chemical cues learned during their respective pre-imaginal stages to find hosts [17, 19]. Early experience of olfactory stimuli associated with their host is an important driver of parasitoid foraging choices, notably leading to host selection [27].

Successful host detection and choice may depend on a parasitoid’s ability to learn different cues during its contact period with its host [2, 28, 29], and is most likely to affect the fitness of the offspring that will develop within the chosen host. Host specificity in parasitoid wasps varies from highly specific, with many species only parasitizing one unique host species, to generalist, with species using a wide range of hosts [30-32]. Furthermore, studies have shown that in certain conditions, specialist parasitoids may successfully lay eggs in new host species [33, 34], which may lead to evolutionary changes in host specificity. Such change in host-specificity may occur through two different strategies. A ‘host switch’ is a sudden or accidental colonization of a new host species by a few individuals capable to establish a new and viable population in the new host, while a ‘host-shift’ is a gradual change of the relative role of a particular host species as primary versus secondary host. Host preference is also by no means static, but is characterized by behavioral plasticity that allows parasitoids to switch hosts when their preferred host is unavailable and by learning host cues associated with positive or negative experiences[35-37].

*Wolbachia* is a single bacterial lineage in the alpha-group of the Proteobacteria. The bacterium benefits from strategies, such as thelytoky that increases the number of *Wolbachia*-transmitting females in the host population [38-42]. Similarly, *Wolbachia* can also successfully spread in their host populations by positively affecting various of their host’s life history traits, including fecundity, and survival to pathogens or environmental stresses [43-48].

The parasitoid wasp *T. brassicae* (Westwood) (Hym.: Trichogrammatidae) is an egg parasitoid widely used as a biological control agent of various Lepidoptera pest species [49-53]. As it is often the case in Trichogramma wasps, *T. brassicae* can reproduce by arrhenotokous parthenogenesis, with unfertilized eggs producing viable haploid male offspring [54-56]. However, in certain lineages of *T. brassicae*, only females can emerge from unfertilized eggs [54, 56, 57]. In these particular cases, studies have shown that the wasps are naturally infected by the endosymbiotic bacterium *Wolbachia*, which modifies the reproductive system of the wasps, such that *Wolbachia*-infected individuals reproduce through the tlythokous parthenogenesis instead [43, 57]. As the embryo grows up in host eggs, their pre-imaginal learning abilities are likely to be affected by direct cues from their hosts, but less by chemical cues linked to host food or environmental chemical cues. We hypothesized that the manipulated pre-imaginal learning and host-preference in *T. brassicae* by *Wolbachia* may support host shift and adaptation to new intracellular environments in parasitoid wasps.

## Material and methods

### Parasitoids

We compared two lineages of *Trichogramma brassicae*: one *Wolbachia* infected (*Wolbachia wBaT.bra* registered as FJ441291 in Genbank), and one uninfected. Previous studies have shown that the *Wolbachia-*infected and *Wolbachia*-free lineages carry the same genetic background [44]. This characteristic allowed us to avoid the use of antibiotics against *Wolbachia*, and their potential confounding effects on the physiology and behavior of antibiotic treated insects [58-60]. Both lineages came from colonies maintained by the Ecology and Behavior Laboratory of the University of Tehran, Iran, and were originally collected in 2016 from a Cornfield in north of Iran (Baboulsar Region, South of Caspian Sea, Iran). Both original lineages have been reared on *Ostrinia nubilalis* (Lep.: Crambidae) in the laboratory for two generations before experimental design. Parasitoids were reared on egg cards (2×5 cm), loaded with one-day old eggs of *O. nubilalis*. The egg cards were placed into emergence canisters and held in incubators at 25±1°C, 16L: 8 D and 50±5% RH. Emergence canisters were closed cardboard cylinders (500 ml, 63×161 mm) with a glass vial (50 ml, 26×93 mm).

For all experiments described below, we used two different Lepidoptera species as host to the parasitoids: the flour moth *Ephestia kuehniella* Zeller (Lepidoptera: Pyralidae) and the grain moth *Sitotroga cerealella* (Lep.: Gelechiidae). Lepidoptera eggs were obtained from a culture maintained at the Insectary and Quarantine Facility, University of Tehran. The cultures were reared at 25±1°C on wheat flour and yeast (5%). Newly emerged and mated female moths were kept in glass containers (500 ml) to provide eggs. Eggs were collected daily to ensure that the eggs used in the experiments were no more than 24 h old. The parasitoids were reared at 25±1°C, 50±5% RH, and 16:8 L: D on eggs of either Lepidoptera hosts for more than ten generations.

### *Wolbachia* screening

We determined *Wolbachia* presence in female wasps, which produced only female offspring [43], by PCR screening for the *Wolbachia surface protein* (*wsp*) gene using the *Wolbachia* specific primers 81F/691R [43,61]. The *Wolbachia* infected line was monitored for the infection throughout the study by PCR. Also uninfected strain was screened randomly to check on any contamination or horizontal transfer between the lines throughout the experiment.

### Experimental design

All treatments were carried out in an insectary room under controlled conditions with a temperature of 25±1°C and 55±5% RH. Individual *Trichogramma* females were presented with patches of host eggs fixed with water onto small pieces of white cardboard. The experiments ended once the female wasps left the patches of eggs that were offered to them. All parasitized eggs were incubated at 25 ± 1 °C, L16:D8, and 50 +/-5% R.H. for 4–5 days until the eggs blackened, suggesting that the parasitoid had pupated within its host. Parasitized host eggs were then cut out of the cardboard patch and placed individually in gelatin capsules to await the emergence of the parasitoid progeny. We then measured the total number of parasitoids that emerged. Parasitism rate was calculated as the ratio of blacked eggs compared to the total number of provided host eggs for each wasp.

Immediately after wasp emergence, mated female wasps were removed from their gelatin capsules and maintained for an hour in new tubes individually. During this period, the wasps were never in contact with hosts, and were kept in a separate growth chamber than their future hosts. The emerging parasitoids sex was determined by morphological differentiation of the antennae [52]. We calculated the sex ratio of the emerging progeny for each female wasp.

#### A) Innate host preference

We first tested the innate preference of newly collected female wasps, infected or not by *Wolbachia*, toward both experimental hosts (*E. kuehniella* and *S. cerealella)*, after emerging from the European corn borer, *O. nubilalis*. Thirty newly emerged wasps infected by *Wolbachia*, and 30 newly emerged uninfected wasps were exposed to eggs from each experimental host in a choice and non-choice test.

In the choice experiment: wasps were exposed to 50 one-day old eggs of *E. kuehniella* and 50 on day old eggs of *S. cerealella* (Ntotal=100 eggs), simultaneously. Eggs of each host species were attached to small cards (1× 5 cm) separately and were placed in a glass tubes (2×10cm).

In the non-choice experiment: newly collected wasps were exposed to patch containing 100 one-day old eggs of each experimental host separately. Each egg card contained 100 eggs fixed on labeled, white cardboard cards (2.5× 5 cm) of either host later exposed to a wasp in a glass tubes (2×10cm). For each test, a naive (no previous oviposition experience) one-day old female wasp was released on each egg patch.

#### B) Host preference after early imaginal experience

To study host preference after early imaginal experience, wasps were reared on either of the two experimental hosts for 10 consecutive generations prior to the experiment. We studied the effect of *Wolbachia* presence and experimental host species during immature developmental time on pre-imaginal learning ability using a full factorial design: each of the two parasitoid lineages were offered to parasitize each of the two Lepidoptera host species, with 30 replicates for each treatment. The four treatments scheme includes Treatment T1: uninfected wasps reared on *E. kuehniella*, T2: uninfected wasps reared on *S. cerealella*, T3: *Wolbachia*-infected wasps reared on *E. kuehniella*, and T4: *Wolbachia*-infected wasps reared on *S. cerealella* (Figure 1).

**Figure 1.**
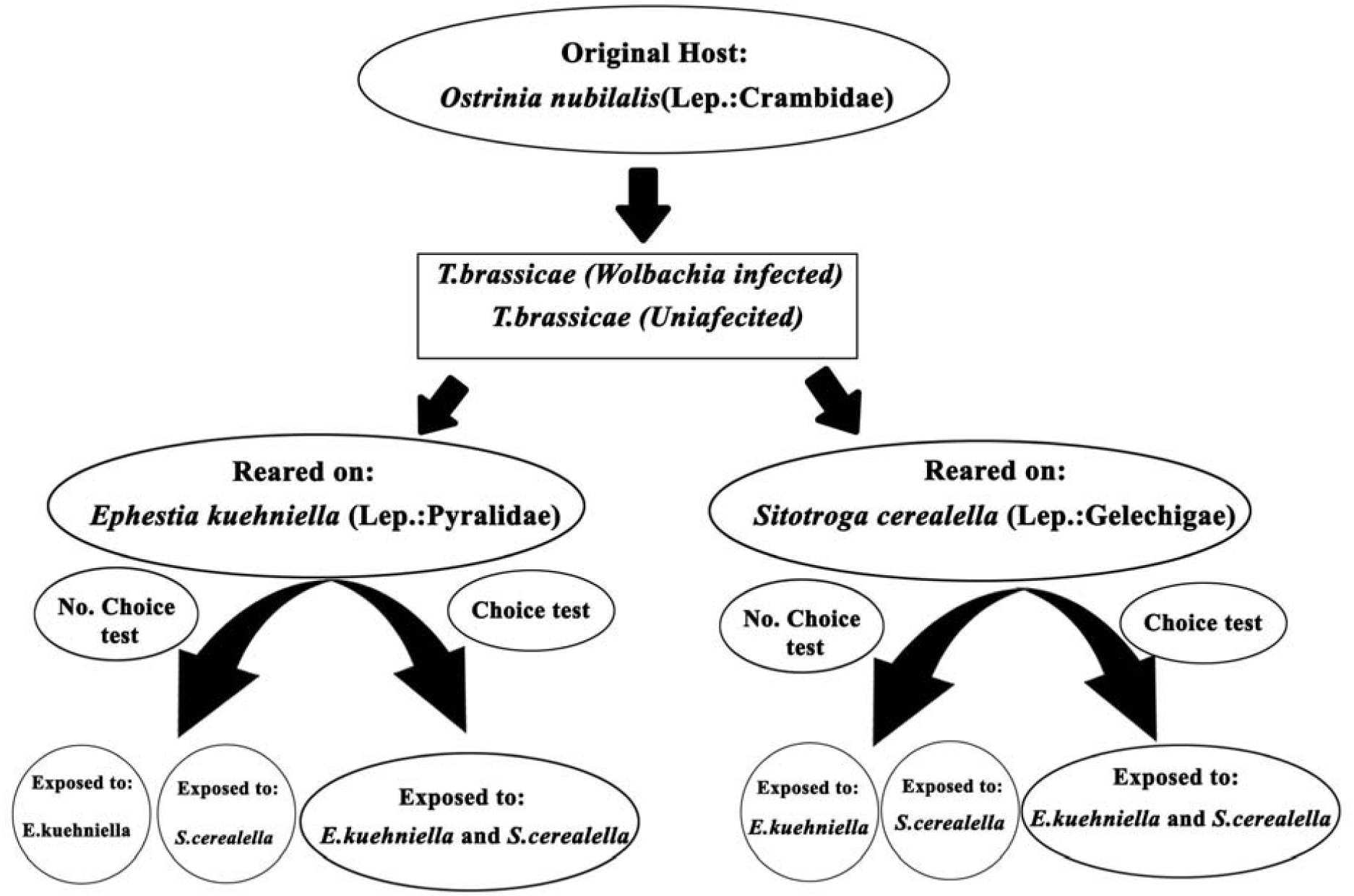
Schematic view of experimental design of host preferences after pre-imaginal learning.

**Figure 2.**
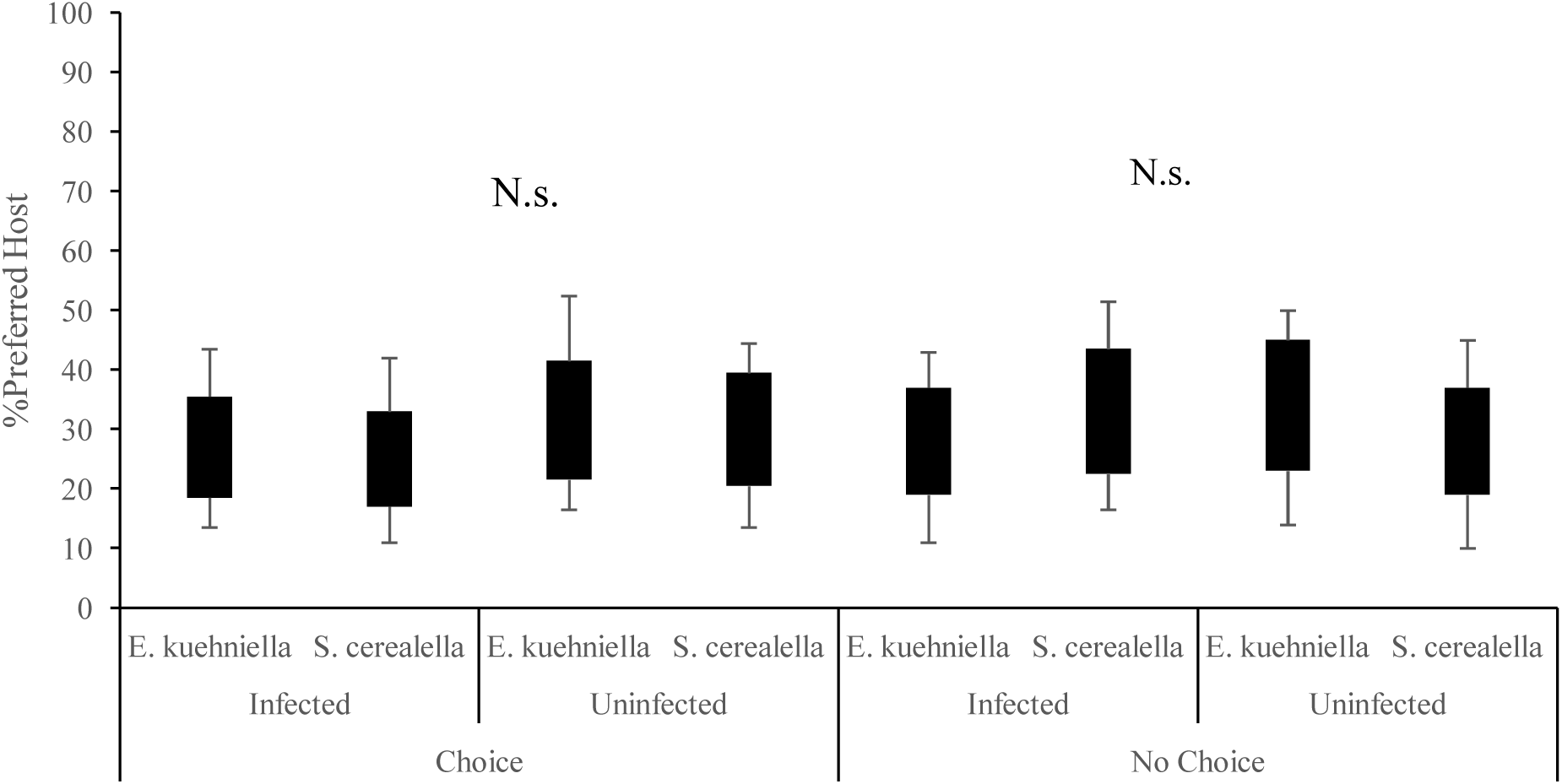
Innate preference of *Trichogramma brassiacae* wasps, *Wolbachia* infected and uninfected, toward *Ephestia kuehniella* and *Sitotroga cerealella* in choice and no choice tests. Wasps were originally all reared on *Ostrinia nubilalis.* (N.s.) stands for Not significant.

### Data analysis

All statistical analyzes were done by SAS software [62]. To compare egg-laying preference of the female wasps, we implemented a Generalized Linear model in the procedure GENMOD of the SAS software (ver. 9.1), with the binomial family error and logit link. After this global test, the least square estimates of the proportions in each level were compared by the Chi-square approximation (an option offered by GENMOD).

The rate of emergence was calculated based on the total number of emerged wasps and the total number of oviposited eggs, and was analyzed for each experiment using generalized linear models based on a binomial logit distribution [63]. The numbers of female and male offspring produced by uninfected wasps were compared by Student’s t-test.

In all cases, the explanatory variables were the host species and the *Wolbachia* infection, and when a significant effect of the treatment was found, the tests were followed by Bonferroni’s *posthoc* multiple comparison tests. The two-by-two comparisons were evaluated at the Bonferroni-corrected significance level of *P* = 0.05/k, where k is the number of comparisons. Data are presented as means ±SE (through the results and Table 1).

**Table 1.** Effects of lineage, host species and the interaction of these two factors on host preference of uninfected and infected wasps. Significant results are shown in bold.

|  |  | lineage |  | Host |  | Interaction |  |
| --- | --- | --- | --- | --- | --- | --- | --- |
| | | $\chi^2$ | P value | $\chi^2$ | P value | $\chi^2$ | P value |
| Innate preference | Choice | 0.43 | 0.511 | 0.02 | 0.889 | 0.02 | 0.889 |
|  | No choice | 0.02 | 0.889 | 0.02 | 0.889 | 0.79 | 0.373 |
| reared on *E. keueneilla* | choice | 4.36 | **0.036** | 1.63 | 0.20 | **7.7** | **0.005** |
|  | No choice | 4.95 | **0.026** | 1.02 | 0.313 | 2.57 | 0.109 |
| Reared on *S. cerealella* | Choice | 14.48 | **0.0001** | 0.24 | 0.622 | 11.57 | **0.0007** |
|  | No choice | 8.93 | **0.002** | 0.61 | 0.433 | 3.76 | 0.052 |

## Results

### Host preference experiments

#### A) Innate host preference

Effects of *Wolbachia* infection status, host species, and their interaction on the preference of the female parasitoid wasps are shown in Table 1. As showed in Figure 1, both lineages displayed no significant preference for either experimental host *E. kuehniella* or *S. cerealella*, and showed the same response toward the two hosts in both the choice and the no choice tests.

#### B) Host preference after early imaginal experience

In the choice experiment: A significant interaction between wasp *Wolbachia* infection status, and host species was observed on the wasps’ host preference (Table 1). The uninfected wasps reared on *E. keueneilla* significantly preferred the hosts that were reared on, *E. keuene*, (χ^2^=9.77, P=0.0018, N=100), while *Wolbachia*-infected wasps did not show any preference for either host (χ^2^=0.26, P=0.611, N=100) (Figure 3). Similarly, *w*hen given the choice, uninfected wasps reared on *S. cerealella* showed a significant preference toward the host they were reared on (χ^2^=14.55, P=0.0001, N=100) (Figure 3), while *Wolbachia* infected wasps reared on *S. cerealella* showed no preference toward either hosts (χ^2^=0.11, P=0.74, N=100).

**Figure 3.**
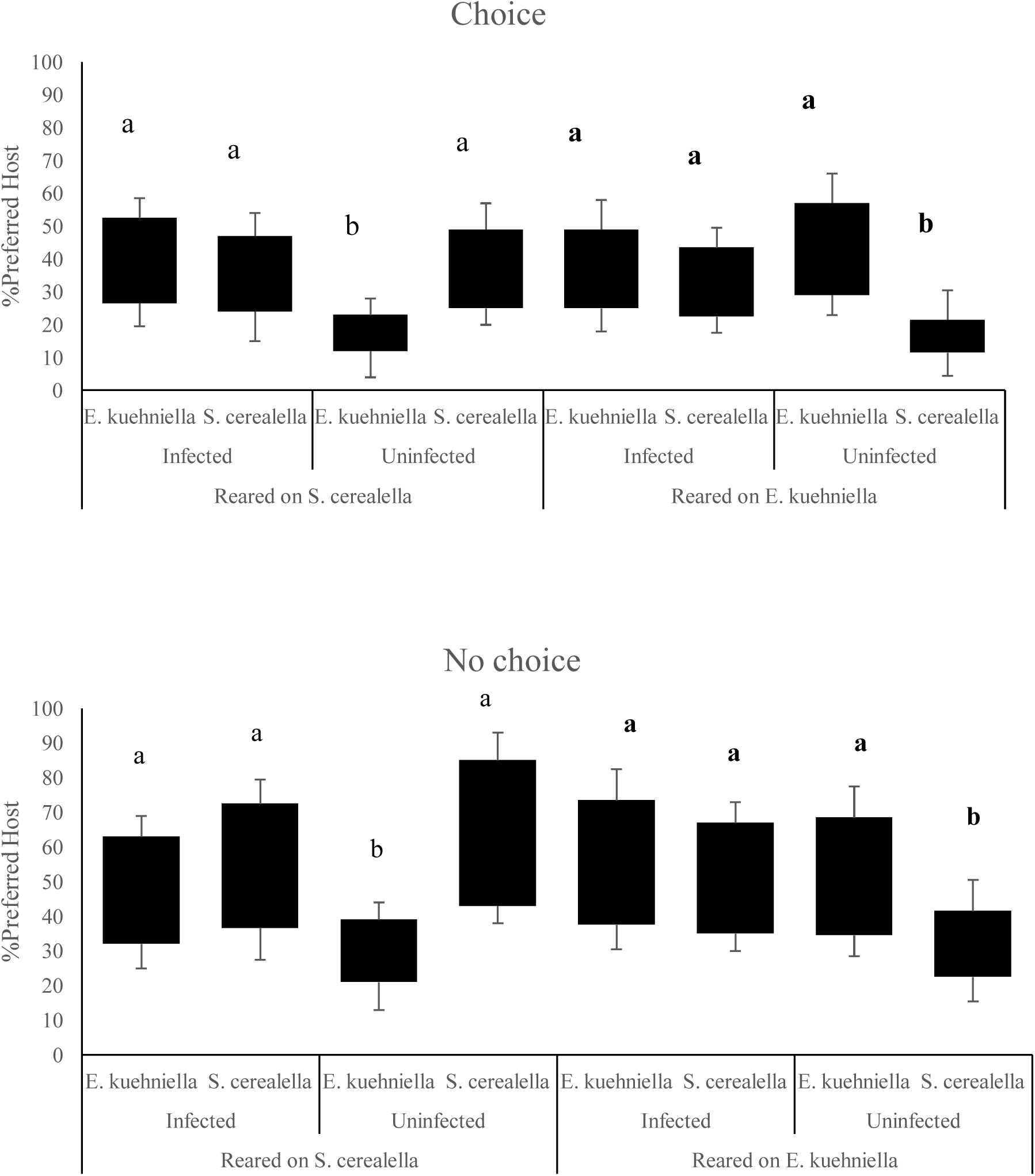
Host preference of *Trichogramma brassiacae* wasps, *Wolbachia* infected and uninfected, after pre-imaginal experience on *Ephestia kuehniella* and *Sitotroga cerealella* in choice and no choice tests. Different letters indicate significant differences.

In the no-choice experiment: Uninfected wasps reared on *E. keueneilla* laid more eggs on *E. keueneilla* (χ^2^=6.79, P=0.009, N=100). However, no significant difference was observed in number of eggs laid in both hosts by infected wasps (χ^2^=0.2, P=0.653, N=100) (Fig 3). Uninfected wasps reared on *S. cerealella*, again significantly oviposited more on the host they were on (χ^2^=11.09, P=0.0009, N=100), while *Wolbachia* infected wasps reared on *S. cerealella* oviposited similarly on both hosts in no choice tests (χ^2^=0.57, P=0.45, N=100) (Figure 3).

### Emergence rate

The parasitoid infection status (χ^2^=8.44, P=0.003), the host species (χ^2^=4.86, P=0.0275), and their interaction (χ^2^=42.41, P<0.0001) had a significant effect on the emergence rate of the parasitoid. Uninfected wasps reared on *S. cerealella* showed significant lower emergence rate when they were reared for one generation on *E. kuehniella* (Figure 4). Similarly, emergence rate of uninfected wasps reared on *E. kuehniella* was significantly lower when wasps were reared on *S. cerealella* for one generation (Figure 4). All other treatments showed similar emergence rates.

**Figure 4.**
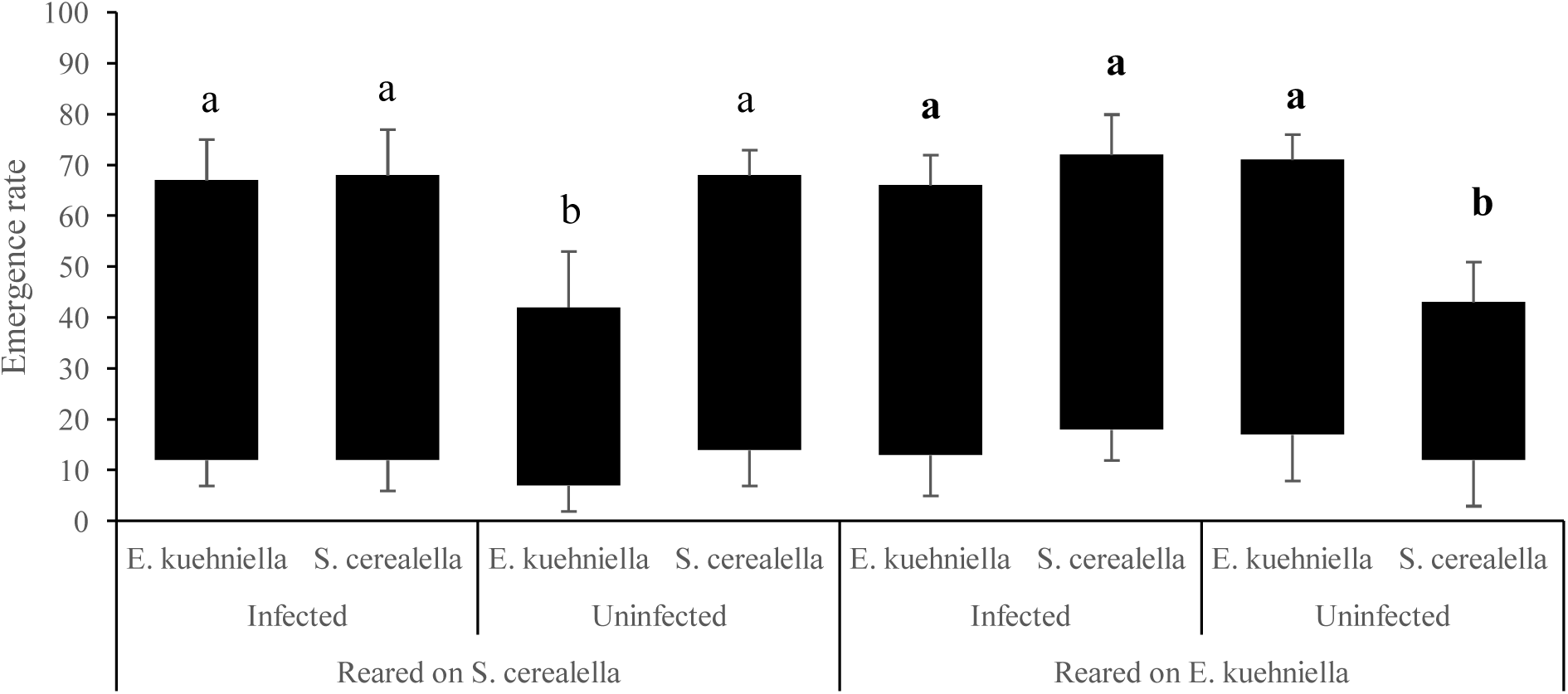
Emergence rate from *Ephestia kuehniella and Sitotroga cerealella* hosts of *Trichogramma brassiacae* parasitoid wasps, either *Wolbachia* infected and uninfected. Different letters indicate significant differences.

### Sex ratio

Reared uninfected wasps on *E. kuehniella* laid more female eggs in *E. kuehniella* eggs (t = 7.668, df = 1, P<0.0001) compared to uninfected wasps exposed to *S. cerealella*; their observed offspring sex ratio was 1:3 (male: female). Also uninfected wasps reared on *S. cerealella* produced more female offspring in patches containing *S. cerealella* eggs (t = 2.677, df = 1, P = 0.015) compared to uninfected wasps exposed to *E.kuehniella*, with a ratio of 1:3. All produced offspring by infected wasps in both hosts were females. Due to *Wolbachia*-induced thelytoky, all offspring from *Wolbachia*-infected wasps were females.

## Discussion

According to our results, *Wolbachia*-infected specimens of the parasitoid wasp *Trichogramma brassicae* do not follow the Hopkin’s host-preference principle, while their uninfected counterparts do. *Wolbachia* manipulates the host preference of the parasitoid wasp, such that female wasps do no discriminate between eggs from *E. kuehniella* and *S. cerealella*. In contrast, uninfected wasps of the same species show a significant preference towards parasitizing the Lepidopteran host species from which they emerged. We also show that when choice is not given, *Wolbachia*-infected wasps lay the same amount of eggs in both Lepidopteran hosts offered, with similar emergence rates from both hosts, contrasting with uninfected wasps, which significantly lay more eggs and consequently show higher emergence rates in the host species they emerged from. The family Trichogrammatidae is one of the earliest branching families of the superfamily Chalcidoidea [64, 65]. Although many members of this family are generalists, host specificity has also been reported [66-68]. The two *T. brassicae* lineages used for this study were collected from a same unique location in Iran, from a same host species, *Ostrinia nubilalis*, and were later shown to share the same genetic background, we believe they originally share similar host preference. Thus, the differences observed in our study are more likely only due to the insects’ *Wolbachia* infection status.

By relaxing strict host preference through modifying pre-imaginal learning ability in its host, generalist *Wolbachia* infected individuals could oviposit in and thus benefit from a wider range of hosts resource [69, 70]. They may as well benefit from a more nutritionally balanced diet, and show higher capacity to confront variable environments [71-73]. In contrast, the information-processing hypothesis suggests that highly efficient decision-making should evolve faster in specialist foragers compared to generalists [74-77]. Additionally, specialists may also evolve as better competitors on their unique hosts, and thus show higher survival, but could also be more susceptible to population fluctuation of the host than generalist species [78, 79].

According to our results, *Wolbachia* infected wasps show similar host preference behavior even after emerging from different hosts. Previously, [80] shown that there are no differences in olfactory responses between conditioned and unconditioned *Wolbachia* infected *T. brassicae* toward peppermint odor. The fixed behaviors were suggested to be a consequence of *Wolbachia* affecting organs structure and function [81, 82], and thus potentially leading to poor information integration, and no differentiation between different host species and host qualities [44]. Studies have also shown that *Wolbachia* can directly affect a range of neurotransmitters with potential impacts on subsequent fitness-related behaviors in various insects [48, 83, 84]. Learning capacity may allow the host to tune behavior to an adaptive solution, especially when the environment is not informative enough for the specific foraging behavior to be optimized [85]. Through manipulating the time dedicated to cues learning and/or information processing, *Wolbachia* may thus appear beneficial for the host that can utilize and develop on different host species.

Specialists have long been seen as potential evolutionary dead-ends [86], but more recent research indicates that transition from specialist to generalist phenotype often occur [87]. Switch to a new host however will most likely require genetic, physiologic, and ecological adaptations from the parasitoid wasps [33, 37, 88, 89]. Our study suggests that symbionts like *Wolbachia* can be powerful evolutionary motors in the transition from specialization to generalization [90-92], as their presence may decrease adaptations costs to the new hosts [93,94]. To switch and survive in/on new hosts, the parasitoids must overcome many new ecological and physiological barriers imposed by the new hosts such as overcoming host immune system [95]. *Wolbachia* has been shown to be involved in nutritional provisioning for its hosts, such as providing some elements missing from their host diet or environment [96-99]. In *T. brassicae*, [100] showed that *Wolbachia* may provide for the larval development of their host, as individuals previously reared on hosts of different qualities showed no fitness differences at the adult stage. The details of this potential provision however remain unclear.

### Conclusion

We show here that *Wolbachia* can indeed manipulate its host preference for certain resource, and maintain a generalist parasitism strategy in *T. brassicae*. Although the details of the genetic and physiological background of the behavioral manipulation still remain to be investigated, we believe this phenotype is conserved in nature as it may support the spread of the symbiotic strain in unstable environments, in which specialists may be at higher risk of extinction. *Wolbachia* has been considered as “a generalist in host use” as it is thought to be present in at least 40% of all insects [101]. Although a strict maternal transmittion of the symbiont would lead to co-evolution of the symbiont and its hosts, phylogenetic studies have shown that the phylogenetic trees of *Wolbachia* strains is rarely congruent with those of their insect hosts [102]. These results suggest that *Wolbachia* transfer horizontally between host species more often than previously thought. By restraining its host ability to preferentially parasitize the Lepidoptera host species it emerge from, the *Wolbachia* strain present in T. brassicae may thus increase its chance to transfer horizontal and potentially establish in a wider range of naïve species populations.

## Acknowledgements

We appreciate Prof. Jean-Sébastien Pierre for his help and comments in improving statistical analyses. Also we are grateful to Seyedeh Samira Qazaei for the technical support provided. This study was financially supported by the University of Tehran, and Academy of Finland (Grant #321543 to AD), and the Marie-Curie Sklodowska Individual fellowship (#120586, Host Sweet Home to AD), but the sponsor had no involvement in the study design, the collection, analysis, and interpretation of data, the writing, or where to submit the paper for publication.

